# High-dimensional Bayesian phenotype classification and model selection using genomic predictors

**DOI:** 10.1101/778472

**Authors:** Daniel F. Linder, Viral Panchal

## Abstract

**Motivation:** In this paper we describe a Bayesian hierarchical model termed ‘PMMLogit’ for classification and model selection in high-dimensional settings with binary phenotypes as outcomes. Posterior computation in the logistic model is known to be computationally demanding due to its non-conjugacy with common priors. We combine a Polya-Gamma based data augmentation strategy and use recent results on Markov chain Monte-Carlo (MCMC) techniques to develop an efficient and exact sampling strategy for the posterior computation. We use the resulting MCMC chain for model selection and choose the best combination(s) of genomic variables via posterior model probabilities. Further, a Bayesian model averaging (BMA) approach using the posterior mean, which averages across visited models, is shown to give superior prediction of phenotypes given genomic measurements.

**Results:** Using simulation studies, we compared the performance of the proposed method with other popular methods. Simulation results show that the proposed method is quite effective in selecting the true model and has better estimation and prediction accuracy than other methods. These observations are consistent with theoretical results that have been developed in the statistics literature on optimality for this class of priors. Application to two well-known datasets on colon cancer and leukemia identified genes that have been previously reported in the clinical literature to be related to the disease outcomes.

**Availability:** Source code is publicly available on GitHub at https://github.com/v-panchal/PMML.

**Supplementary information:** Supplementary data are available online.

## 1 Introduction

The Bayesian paradigm offers a natural statistical framework for model and variable selection, namely by assigning priors that convey a prior belief in model sparsity. Such priors then lead to sparsity *a posteriori*; i.e., the data modifies our prior beliefs about the importance of variables or groups of variables, which is expressed through the posterior distribution. The additional assumption of model sparsity (that many parameters are zero), either through prior distribution or penalty function, is actually necessary in some situations, like for instance in high-dimensions. Otherwise the inferential problems may not be well posed or identifiable. Although there are various situations that are deemed to fall under the umbrella term of high-dimension, our focus is on data where sample sizes are small relative to the number of predictors, *p* > *n*. To overcome the associated identifiability issues when *p* > *n*, the frequentists have approached the problem by adding penalties to classical objective functions (Breiman, 1995; Tibshirani, 1996; Golub *et al*., 1979; Hoerl and Kennard, 2000; Fan and Li, 2001; Efron *et al*., 2004; Zou, 2006; Van de Geer, 2008; Zhang and Huang, 2008).

Many approaches have been developed for variable and model selection in high dimensional problems. The LASSO, proposed by Tibshirani (1996), is perhaps the most well known, with many extensions and adaptations since. Wu *et al*. (2009) and Ravikumar *et al*. (2010) extended the LASSO to logistic regression in high-dimensional settings, some closely related approaches include smoothly clipped absolute deviation (SCAD) (Fan and Li, 2001), Dantzig selector (Candes and Tao, 2007), the adaptive LASSO (Zou, 2006), and the elastic net (Zou and Hastie, 2005). More recently, Fan and Lv (2008) proposed the sure independence screening based method to perform variable selection. Their approach involves two steps: (1) reduce the dimension from high to one smaller than the sample size by performing correlation screening, and (2) improve variable selection by a penalized likelihood approaches such as SCAD, the Dantzig selector, LASSO or adaptive LASSO.

Bayesian versions of many of these frequentist approaches have been developed where the posterior modes match the corresponding frequentist solution (Park and Casella, 2008; Zellner, 1976; Li and Lin, 2010; Leng *et al*., 2014; Linder *et al*., 2016), these are sometimes referred to as shrinkage priors. Other work using shrinkage priors has been developed in the variable selection literature (Bae and Mallick, 2004; Griffin and Brown, 2005). Most of these previously mentioned priors are what (Johnson and Rossell, 2012) call local priors, and they have pointed the serious limitation that local priors give models lacking the stong model selection consistency criteria of (Johnson and Rossell, 2012). This implies that even in the presence of ever increasing amounts of data, the posterior probability of the true model will be set to zero. Further, some priors from this class have also been shown to lead to sub-optimal estimation rates. To address the first limitation, two new classes of non-local priors were proposed by Johnson and Rossell (2010) and studied in detail for Bayesian model selection in high-dimensional settings in (Johnson and Rossell, 2012) and (Rossell *et al*., 2013). Models using these non-local priors achieve strong model selection consistency, and Nikooienejad *et al*. (2016) has adapted these non-local priors to Bayesian variable selection with binary outcomes. Two challenges associated with their non-local priors are: (1) exact MCMC is intractable, and thus posterior computation requires an approximate MCMC using the Laplace approximation; (2) to our knowledge the posterior contraction and estimation rates associated with non-local priors are unknown. This is in contrast to the point mass mixture priors that our model uses. It has been shown, Atchadé *et al*. (2017), that this form of prior leads to optimal posterior contraction rates in the logistic model.

Another approach to Bayesian variable and model selection uses so-called two-group priors. Two-group models use a mixture of two distributions for the prior: one for the zero components and another for the non-zero components. The resulting posterior then attempts to separate zero (noise) and non-zero (signal) components through this two-group mixture structure. Popular two-group approaches include mixture-of-normals approximation to spike-and-slab priors (George and McCulloch, 1997), stochastic search variable selection (SSVS) (George and McCulloch, 1993), and MCMC based stochastic variable selection (Lee *et al*., 2003). Besides their intuitive nature, there is strong theoretical motivation for two-group priors due to their optimality, see Johnstone and Silverman (2004); Castillo *et al*. (2012). Because of this the two group priors are often considered to be a Bayesian gold standard. Unfortunately, the associated MCMC in the two-group models can be challenging. This is primarily because the MCMC procedures must explore large model spaces in a discrete way George and McCulloch (1993); Green (1995); Carlin and Chib (1995); Kuo and Mallick (1998). When the proposal distributions are not carefully tuned, which is an important ingredient for procedures like the widely used Metropolis-Hastings algorithm, the resulting MCMC chain will have poor mixing and the inferences based on them may be useless. This is in contrast to many of the continuous shrinkage priors, where block updating of parameters via Gibbs sampling can lead to highly efficient MCMC strategies.

The main contribution of this paper is a Bayesian hierarchical model with an efficient MCMC strategy, and its software implementation, for relating a possibly high dimensional set of genomic predictors with a binary phenotype. Our computational strategy uses a data augmentation result developed in Polson *et al*. (2013), coupled with measure-theoretic results in Gottardo and Raftery (2008), to perform efficient and exact MCMC in the logistic model with point mass mixture priors. We show that the posterior gives superior model selection and predictive properties in simulation and real data examples. Beyond the current section, the paper is organized as follows. In Section 2, we describe the Bayesian hierarchical model and detail how data augmentation of the logistic likelihood can be used in combination with the prior for efficient posterior computation via Gibbs sampling. We compare the proposed model with current state of the art methods in terms of variable selection, estimation, and prediction in Section 3 through simulation and on real data examples. We conclude with discussion and future work in Section 4.

## 2 Methods

### 2.1 Bayesian hierarchical model

### 2.2 Pólya-Gamma likelihood representation

Consider data that is of the form *D* = ((*y*_*i*_, *n*_*i*_), *x*_*i*1_, …, *x*_*ip*_) for *i* = 1, …, *N*, where *y*_*i*_ denotes the number of successes out of *n*_*i*_ trials, and *x*_*ik*_ is the *k*^*th*^ covariate value of subject *i*. A binomial assumption on the data combined with a linear model for the log odds of success gives a likelihood function, written as a function of *β*, as

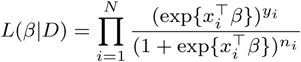

Typically *n*_*i*_ = 1, so that each *y*_*i*_ is either 0 or 1, which in the current context is one of the two possible phenotypes; i.e., good prognosis vs. poor prognosis, or malignant vs. benign. This is the standard logistic regression model, and the covariate *β*_*k*_ determines the effect of the *k*^*th*^ predictor on the probability that *y* is of type 1. For the remainder of the paper, we will assume the feature vector has been augmented with a 1 in the first position and the coefficient vector *β* continas the intercept term *β*_0_ in the first position.

Bayesian data analysis in the logistic model has until recently been computationally difficult, in contrast to the probit model, where data augmentation strategies have been available for some time, Chib (1995). The appeal of the logistic model over the probit model is its straightforward interpretation of covariate effects on the response in terms of odds ratios, and is thus usually preferred in epidemiological and clinical research to probit regression. Efforts to implement similar augmentation strategies in the logistic model have been less fruitful, and have either relied on approximations or require multiple levels of latent variables. Motivated by these challenges, Polson *et al*. (2013) developed a data-augmentation scheme for Bayesian logistic regression that is exact and requires just a single layer of latent variables. The representation has the form

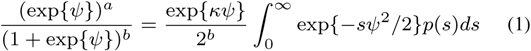

where *κ* = *a* − *b/*2 and *p*(*s*) is the density function of a Pólya-Gamma random variable, *S ∼ PG*(*b*, 0). In general, a Pólya-Gamma random variable *S* with parameters *b* and *c* is distributed as

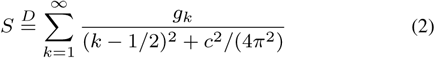

where *g*_*k*_ ∼ *G*(*b*, 1) are independent gamma variates. The corresponding density function of a *PG*(*b, c*) random variable is of the form

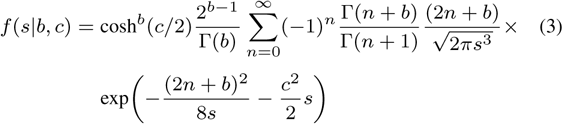

The integrand in (1) is proportional to a normal density (in *ψ*), and thus conditional on *s, ψ*|*s* is also normal. Polson *et al*. (2013) have also shown that the full conditionals, *s*|*ψ* ia also Pólya-Gamma and have developed an efficient sampler for this. It is this representation in combination with a point mass mixture prior that allows us to formulate our scalable MCMC algorithm presented later.

### 2.3 Variable selection priors

A point mass mixture prior on the regression parameters consists of a point mass at 0 mixed with a continuous distribution, with mixing weight *ω* ∈ [0, 1]. More formally, consider the Dirac mass function *δ*_0_, which is concentrated at 0, and a continuous distribution *F* with support on the real line. Further, assume *F* is absolutely continuous with respect to Lebesgue measure *λ* and hence admits density function *f*. The corresponding mixture measure, Π = (1 *− ω*)*δ*_0_ + *ωF*, has density function (Radon-Nikodym derivative)

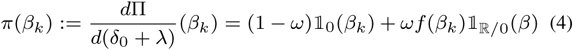

see Gottardo and Raftery (2008). The above mixture is a two-group assumption that *β*_*k*_ is either 0 with probability 1 *− ω*, or non-zero and distributed according to *f* with probability *ω*. Heavy-tailed density functions for *f*, like the double exponential (i.e., Laplace), have been shown to lead to the optimal posterior properties mentioned in the introduction Johnstone and Silverman (2004); Atchadé *et al*. (2017). Our proposed Bayesian hierarchical model is

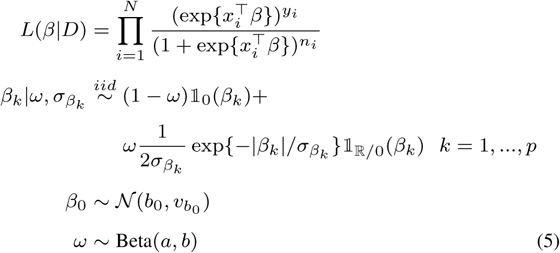

The intercept term is separate from the point mass mixture hierarchy on the other coefficients because it is does not correspond to any genomic predictor, and it should not influence, at least explicitly, the probabilities that genomic features are important. We do this by placing a separate normal prior on the intecept, and eventually we marginalize it away from the posterior. The beta prior on mixture weight, *ω*, specifies a distribution on the probability that a genomic feature is important; i.e., non-zero parameter, in which case it comes from the continuous component and otherwise is set to zero by the indicator function. The parameter *ω* also controls the overall model size through the hyperparameters *a* and *b*. Consider a model ℳ_*k*_, with a configuration of *k* non-zero parameters and *p* − *k* zero parameters, then *p*(ℳ_*k*_|*ω*) = *ω*^*k*^(1 *− ω*)^*p−k*^ follows from the prior on regression coefficients in the above hierarchy. Then the induced prior on the model space has density for model as *ℳ*_*k*_ as 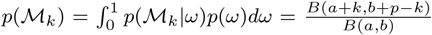, see Castillo *et al*. (2015) and Nikooienejad *et al*. (2016). Here a small *a* and large *b* gives greater probability to sparse models and low probability to larger models. We set 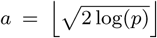 which is the universal thresholding rule as our best guess *a priori* for the number of non-zero signals, and we set *b* = *p − a* to represent the remaining number of superfluous (noise) variables. The hyperparameters 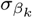 are the variance components for non-zero parameters. Finally, we set *b*_0_ to 0 and variance components 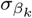 and 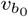 to 10^3^ which produces relatively flat priors on the intercept and slab. We have observed emprically that analysis is robust to choices of these variance components with similar qualitative results within ranges 10^0^ to 10^5^ for these.

### 2.4 Posterior computation

Although priors of the type in Equation (4) have been studied extensively in the literature, our focus in this paper is to use results presented in Gottardo and Raftery (2008), in combination with the representation in Equation (1), to provide software implementation of a highly efficient MCMC routine for posterior sampling of the proposed hierarchical model. Posterior computation for this model has proven to be challenging since the logistic likelihood does not lead to conditional conjugacy for model parameters, otherwise a straightforward Gibbs block updating could be carried out. Thus, Metropolis-Hastings (MH) type algorithms have been required in this setting. Although MH proposal distributions can be tuned to give optimal acceptance rates, hierarchical models with the priors in Equation (4) require complicated proposals of the same form; i.e., ones dominated by measures of the form (*δ*_0_ + *λ*)^*p*^, for which tuning is not easy. Inadequate tuning can have serious consequences on the quality of the resulting chains, particularly in terms of mixing.

The hierarchy in Equation (5) does not immediately lead to a straightforward expression for the full conditional distributions of the model parameters; however, using the following mixture representation for continuous (non-zero) component

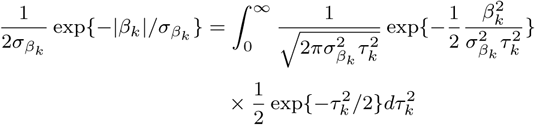

which is the Laplace distribution, leads to analytic full conditionals after data augmentation of the likelihood. The above mixture representation is the one exploited in the original Bayesian lasso of Park and Casella (2008); Andrews and Mallows (1974); West (1987). The data augmentation of the likelihood follows from Equation (1) by introducing *q*_*i*_ for each likelihood component. These representations result in our efficient MCMC routine via Gibbs sampling that we have described in Algorithm 1 below, see Appendix for derivations. Running Algorithm 1 produces a chain with stationary distribution *π*(*β, β*_0_, *ω, τ, q*|*D*), and the target distribution *π*(*β*|*D*) may be computed by marginalizing over (*β*_0_, *ω, τ, q*).

### 2.5 Model selection

We obtain MCMC samples for all model parameters using the above mentioned posterior sampling scheme in Algorithm 1 for the PMMLogit model. There are several ways to perform variable/model selection with samples from the MCMC chain, and we describe the two approaches we consider. Due to the discrete nature of the prior, it is straightforward to select as the final model the one(s) with highest posterior probability(ies). This is because at each pass of the Gibbs algorithm certain coefficients are either in (non-zero) or out (zero) of the model at each iteration. Importantly, the ability to do model selection this way avoids two problems with common approaches: (1) marginalization and (2) ad-hoc cutoffs. Common approaches will often marginalize over each parameter separately, and then perform model selection based on the marginal posterior probabilities or Bayesian credible intervals with continuous priors. Marginalization may miss important joint relationships between model parameters. Further, a model selection process based on marginal posterior probabilities or credible intervals requires setting a thresholding criterion to evaluate the significance of the variables; i.e., by setting those equal to zero if they fall under a thresholding rule. A common rule, which is supported by decision theory arguments, is to use 0.5 as a thresholding rule for both posterior probabilities and credible interval levels. In certain situations this kind of rule may lead to unsatisfactory results. For instance, consider the thresholding value is set to 0.5, and there are two single-variable models, ℳ_1_ and ℳ_2_, where the posterior probability of the single variable in ℳ_1_ is 0.55 and the variable in ℳ_2_ has posterior probability 0.45. Such a rule would discard the variable in the model ℳ_2_, although both variables from ℳ_1_ and ℳ_2_ are of the same relative importance compared to other variables. Adjusting this cutoff, to say 0.25, would increase the expected proportion of variables found to be significant, and include the variable in model ℳ_2_ in this case. In general, this strategy would also increase the number of false positives, leading experimentalists trying to validate these findings on more wild goose chases.

#### Algorithm 1 Posterior computation via Gibbs sampling using the point-mass-mixture prior

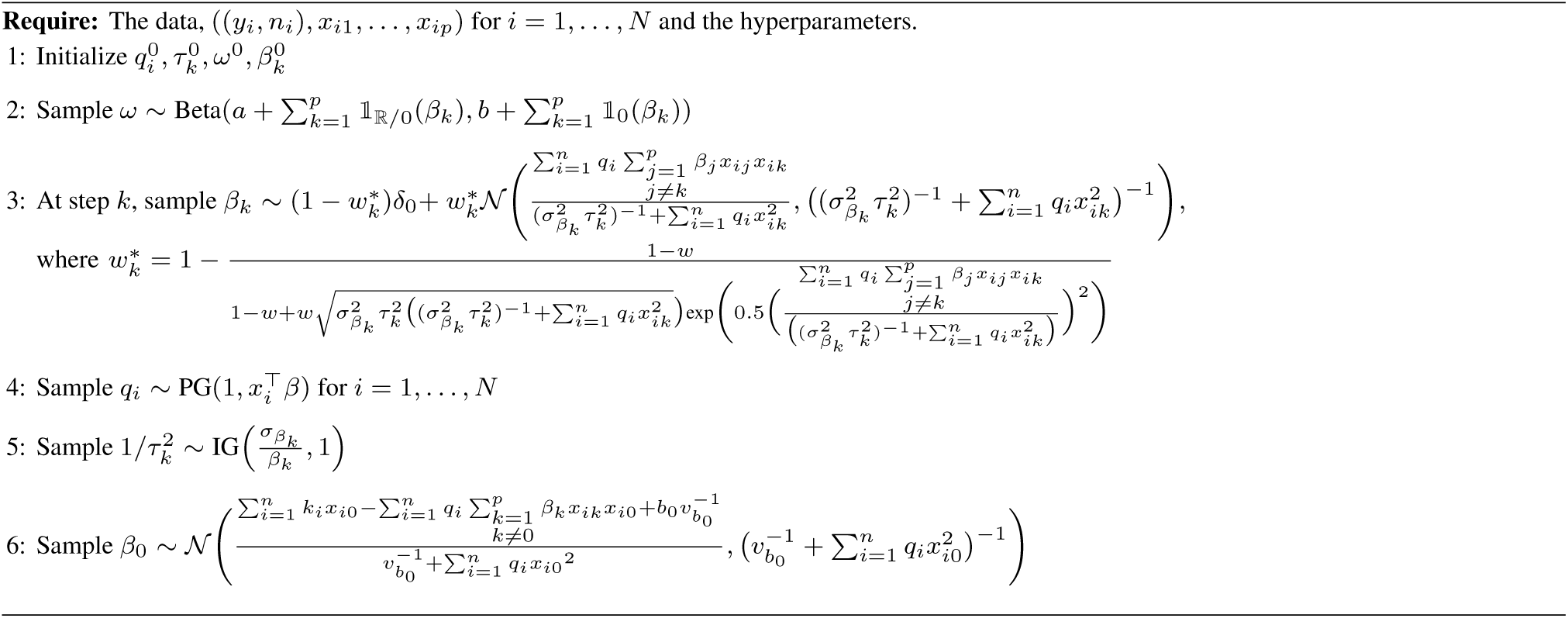

The software package we have devloped for Algorithm 1 produces a list of all visited models that is sorted according to the frequency of each model. We suggest performing model selection by choosing the most frequent model as the final model when its posterior probability clearly separates it from the rest of the competing models. For instance, if ℳ* is the most visited model, with say posterior probability 0.95, with all other models having posterior probabilites *<* 0.01, it would be quite reasonable to select ℳ*. Then, researchers could further investigate the genomic variables occurring in ℳ* for the plausibility of their biological association with the phenotype. Such clear model separation is not always observed, and several competing models may have similar posterior probabilities. Since it is often the researcher’s goal to obtain a list of the most important candidate genes, we describe a clustering approach that combines the information across models. First, for each variable we compute the absolute value of the posterior mean and standard deviation, 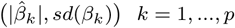, from the MCMC samples. Next, we apply two-means clustering to these *p* 2-vectors to select non-zero variables. The idea behind the procedure is that values near (0, 0) correspond to parameters that were most often zero with little variability, and those far from (0, 0) are variables with potentially large absolute mean; i.e., non-zeros, and/or variation due to within model sampling where it was non-zero, or across model variability.

### 2.6 Prediction

For prediction we implement a simple strategy that is akin to Bayesian model averaging (BMA). We do this by simply using the posterior mean of the coefficients, obtained from the MCMC chain, and use it to make predictions of the outcome given covariate values. The posterior mean of the proposed hierarchical model is a weighted average of the coefficient values over all visited models, and weighted according to each visited model’s posterior probability, or importance. Averaging over models accounts for model uncertainty and leads to superior predictive coverage as compared to the predictive coverage of a single model (Hoeting *et al*., 1999; Carvalho *et al*., 2010). Such strategy also achieves the smallest risk under squared error loss. Although the final model based on the posterior mean may not have exact zeros, it should nonetheless have better predictive performance than using a single model, for example, only the one with highest posterior probability. We demonstrate the predictive performance of our method using this BMA-like strategy in two real data examples.

## 3 Results

### 3.1 Simulation studies

We examine the performance of our proposed method, as well as other methods empirically in this section. Several simulations with different combinations of *p* and model sparsity patterns are considered. We also test different ranges of regression coefficient sizes. The different combinations of model sparsity and parameter magnitude are as given below:

1. *p* ∈ {300, 500, 1000}
2. Number of non-zero coefficients ∈ {5, 10, 15}
3. Non-zero coefficients of *β* ∈ {*β*_1_, *β*_2_, *β*_3_, *β*_4_, *β*_5_, *β* _6_}, where

- *β*_1_ = {1.82, 1.74, 1.78, 1.76, 1.81}
- *β*_2_ = {.82,.74,.78,.76,.81}
- *β*_3_ = {1.82,1.74,1.78,1.76,1.81,1.9,1.85,1.5,1.8,1.6}
- *β*_4_ = {.82,.74,.78,.76,.81,.9,.85,.5,.8,.6}
- *β*_5_ = {1.82,1.74,1.78,1.76,1.81,1.9,1.85,1.5,1.8,1.6,1.84,1.75,1.9,1.8,1.7}
- *β*_6_ = {.82,.74,.78,.76,.81,.9,.85,.5,.8,.6,.84,.75,.9,.8,.7}

We keep the sample size *n* fixed at 300, and for each simulation we generate the design matrix *X* from a multivariate normal distribution with mean 0 and covariance matrix *I*, where *I* is the identity matrix.

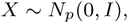

and the response variable *y* is generated as

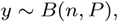

where *n* is the number of observations and *P* = exp(*Xβ*)*/*(1 + exp(*Xβ*)). A total of 18 possible combinations of *p* and *β* is examined. The proposed hierarchy of priors allows user to specify higher weight for relevant predictors. In all simulations, we set weight of the last two non-zero coefficients to be 0.9 to force those to be in the model. Although we specify weights for these two coefficients, we do not consider them in the comparison of performance.

#### 3.1.1 Performance measures

We assess the performance of each method in terms of coefficient estimation, outcome prediction, and variable selection. Specifically, the measures we report for evaluating the performance of each method are:

1. **Frobenius norm**. To evaluate the ability to estimate coefficients, we calculate the Frobenius norm (FN) of the difference (Δ) between the true *β*s and estimated *β*s for each method. FN is calculated as

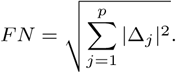
2. **Area under the ROC curve (AUC)**. We use AUC to measure the prediction performance for the binary outcomes of each method. To calculate AUC, we use predicted probabilities with a threshold 0.5 against the observed outcomes. We used the R package ‘ROCR’ (Sing *et al*., 2005) available from CRAN.
3. **True positive rate (TPR)** and **False discovery rate (FDR)**. We quantify the performance of variable selection by finding the average TPR and FDR over 100 iterations. A true positive (TP) is defined as a non-zero variable that is correclty selected as non-zero in the final model, and a false positive (FP) as a zero coefficient that is incorrectly selected as non-zero in the final model. The TPR and FDR are defined as follows:

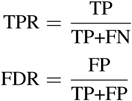

#### 3.1.2 Simulation results

All simulation results are based on 100 replications. The results in Table 1 are for the average Frobenius norm for *β* ∈ {*β*_1_, *β*_3_, *β*_5_}. It can be seen that in all cases, PMMLogit outperforms other methods in terms of estimation. Additionally, as the sparsity level decreases, the difference between PMMLogit and iMOMLogit becomes more pronounced. This was because iMOMLogit leads to too much model sparsity as the number of true coefficients increases, which indicates that PMMLogit is more adaptive to different sparsity levels. Further, the observed superiority in estimation of the coefficients is consistent with the known theoretical optimality for estimation and posterior contraction associated with PMMLogit. The normal prior NP performs the worst among all methods since virtually no sparsity information is incorporated by it. The comparison of the average AUCs for all methods is shown in Table 2 for *β* ∈ {*β*_1_, *β*_3_, *β*_5_}. We again see a similar pattern when comparing PMMLogit to iMOMLogit, for sparse models (*β*_1_) both methods performs similarly, but as the true underlying model becomes more complex, outcome prediction improves significantly for PMMLogit. It is also interesting to see that ISIS and LASSO have better AUCs than iMOMLogit for non-sparse models, with only marginally worse AUCs than PMMLogit. This however comes at a heavy price, in terms of the false discovery rate, for ISIS and LASSO, as we see next.

**Table 1.**
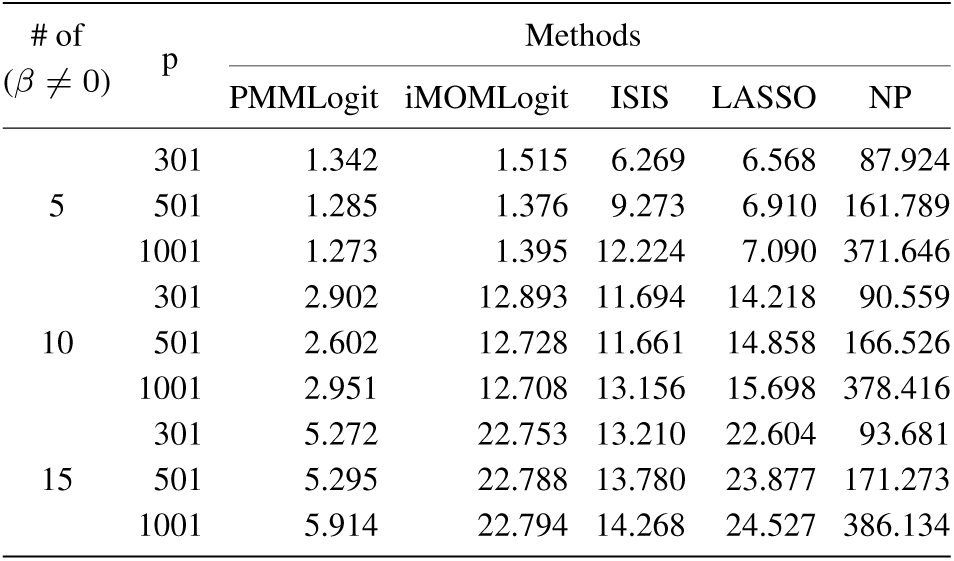
Average Frobenius norm for *β*_1_, *β*_3_ and *β*_5_.

| # of<br>($\beta \neq 0$) | p | Methods | | | | |
| --- | --- | --- | --- | --- | --- | --- |
|  |  | PMMLogit | iMOMLogit | ISIS | LASSO | NP |
| 5 | 301 | 1.342 | 1.515 | 6.269 | 6.568 | 87.924 |
|  | 501 | 1.285 | 1.376 | 9.273 | 6.910 | 161.789 |
|  | 1001 | 1.273 | 1.395 | 12.224 | 7.090 | 371.646 |
| 10 | 301 | 2.902 | 12.893 | 11.694 | 14.218 | 90.559 |
|  | 501 | 2.602 | 12.728 | 11.661 | 14.858 | 166.526 |
|  | 1001 | 2.951 | 12.708 | 13.156 | 15.698 | 378.416 |
| 15 | 301 | 5.272 | 22.753 | 13.210 | 22.604 | 93.681 |
|  | 501 | 5.295 | 22.788 | 13.780 | 23.877 | 171.273 |
|  | 1001 | 5.914 | 22.794 | 14.268 | 24.527 | 386.134 |

**Table 2.**
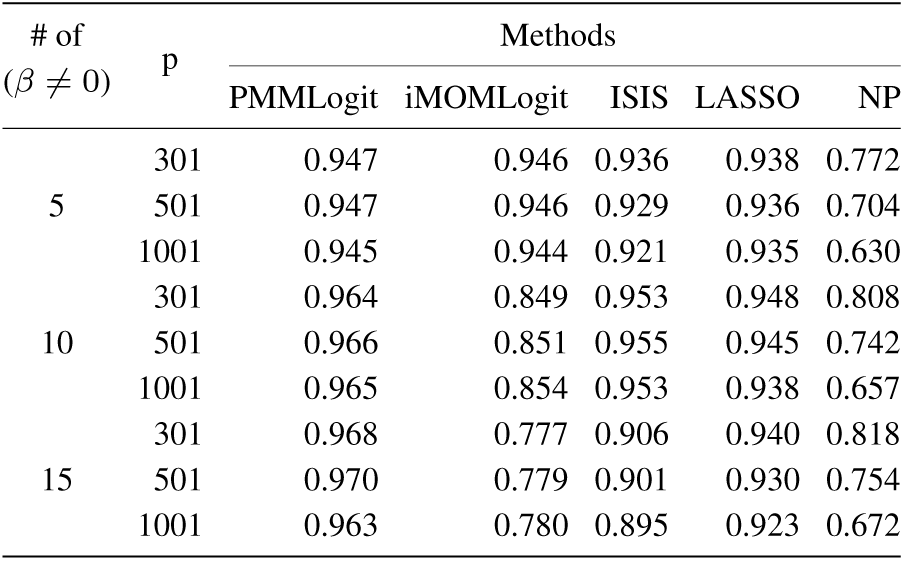
Average AUC for *β*_1_, *β*_3_ and *β*_5_.

| # of<br>($\beta \neq 0$) | p | Methods | | | | |
| --- | --- | --- | --- | --- | --- | --- |
|  |  | PMMLogit | iMOMLogit | ISIS | LASSO | NP |
| 5 | 301 | 0.947 | 0.946 | 0.936 | 0.938 | 0.772 |
|  | 501 | 0.947 | 0.946 | 0.929 | 0.936 | 0.704 |
|  | 1001 | 0.945 | 0.944 | 0.921 | 0.935 | 0.630 |
| 10 | 301 | 0.964 | 0.849 | 0.953 | 0.948 | 0.808 |
|  | 501 | 0.966 | 0.851 | 0.955 | 0.945 | 0.742 |
|  | 1001 | 0.965 | 0.854 | 0.953 | 0.938 | 0.657 |
| 15 | 301 | 0.968 | 0.777 | 0.906 | 0.940 | 0.818 |
|  | 501 | 0.970 | 0.779 | 0.901 | 0.930 | 0.754 |
|  | 1001 | 0.963 | 0.780 | 0.895 | 0.923 | 0.672 |

We report the average TPR and FDR in Figures 1 and 2, respectively. Figure 1 indicates the average TPR for PMMLogit, ISIS, LASSO and NP is much higher than iMOMLogit. On the other hand, the average FDR for PMMLogit and iMOMLogit are nearly identical and very low. ISIS comes in second compared to the two, but still has a much higher FDR, while LASSO and the NP have unacceptably high FDRs. The overall message from this empirical analysis is that PMMLogit demonstrates much better performance in terms of power and signal detection compared to iMOMLogit, and does this with an even lower (although marginally) false discovery rate. The tables and figures for other combinations are provided in supplementary materials and demonstrate similar findings.

**Fig. 1.**
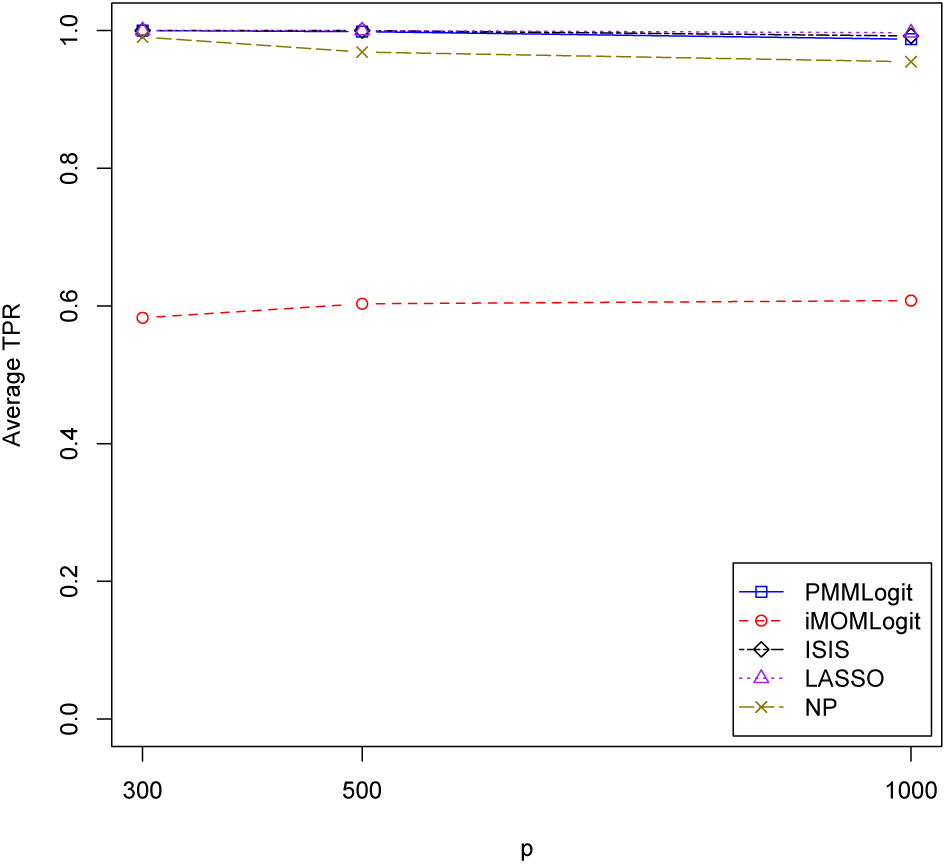
Average true positive rate for *β*_3_.

**Fig. 2.**
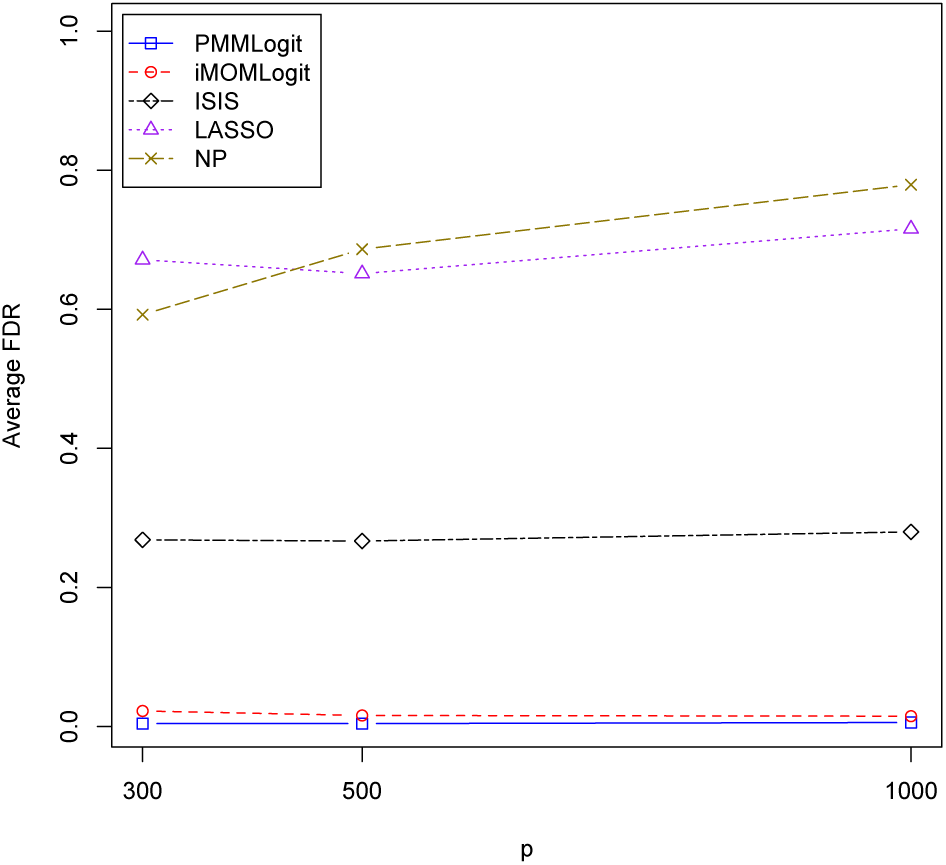
Average false discovery rate for *β*_3_.

### 3.2 Real Data analysis

In this section, we illustrate the utility and performance of PMMLogit on a colon cancer data set and leukemia gene expression data which contain a large number of genomic variables and binary phenotype. The colon cancer data contains 2000 genes and 62 samples. This data set originated from colon adenocarcinoma specimens. The raw colon cancer data contained expression levels of approximately 6,500 genes, and we use a subset of 2000 genes identified by Alon et al. (1999) based on the confidence in the measured expression levels. This data set is publicly available in the R package **plsgenomics**. The second data set is a leukemia data set from the leukemia microarray study of Golub et al., which contains expression levels of 7,129 genes and 72 samples. This data set was calibrated and normalized to obtain robust multiarray average (RMA) expression levels (Irizarry *et al*. (2003)).

#### 3.2.1 Colon cancer data

Alon et al. (1999) analyzed gene expression in 40 tumor and 22 normal colon tissue samples with an Affymetrix oligonucleotide array complementary to more than 6,500 human genes. We use a total of 2000 genes for each colon tissue sample. The goal of our analysis is to identify genes associated with tumor tissue. Three preprocessing steps as described in Dudoit *et al*. (2002) were applied to the design matrix containing gene expression levels. As a result of preprocessing steps, data for 1224 genes across 62 samples remained for further analysis.

We divide the data into training and test sets. The training set contained 43 samples, with 28 tumor and 15 normal samples. The test set contained 19 samples, with 12 tumor and 7 normal samples. To compare the methods, we calculate the error rate for predicting the phenotype in the 19 observations in the test data. We present the results of our method and the others in Table 3. It can be seen that the error rate for both PMMLogit and NP is 5.26%, since they both gave the same number of misclassifications. The error rate for iMOMLogit, ISIS, and LASSO is 10.53%, which is clearly worse than PMMLogit. We report results on the final model selected for each method using the full dataset. Our method selected two genes for the final model named ‘Hsa.36689’ and ‘Hsa.37937’. The genes selected by the method based on the non-local prior included only one gene named ‘Hsa.36689’ while ISIS identified two genes named ‘Hsa.36689’ and ‘Hsa.35496’. The LASSO method and the method based on normal prior resulted in non-sparse models containing 10 and 550 genes, respectively.

**Table 3.** Average error rate and selected genes for the colon cancer data set

| Method | Error rate | Selected genes |
| --- | --- | --- |
| PMMLogit | 5.26 % | Hsa.36689 - Hsa.37937 |
| iMOMLogit | 10.53 % | Hsa.36689 |
| ISIS | 10.53 % | Hsa.36689 - Hsa.35496 |
| LASSO | 10.53 % | 10 genes were selected. |
| NP | 5.26 % | 550 genes were selected. |

*Hsa*.*36689* This gene is described as H. Sapiens mRNA for GCAP-II/uroguanylin precursor. According to Notterdam et al., 2001, uroguanylin and guanylin are significantly reduced in early colon tumors with very low expression. Hsa.36689 has been shown to be significantly associated with colon cancer (Shevade and Keerthi, 2003; Li *et al*., 2002; Jing *et al*., 2010; Garzón and González, 2015; Tan *et al*., 2010; Hossain *et al*., 2013).

*Hsa*.*35496*, also known as H. Sapiens Integrin Alpah-6 Precurson, was identified by the ISIS method. ‘Hsa.35496’ has been reported once by Domany *et al*. (2009) using a coupled two-way clustering approach on the colon cancer data set.

*Hsa*.*37937* Our method also selected this gene in the final model, and it is characterized as ‘Myosin Heavy Chain, Nonmuscle (Gallus gallus)’. It was previously reported to play a role in colon cancer in several studies (Horaira *et al*., 2018; Hossain *et al*., 2013).

#### 3.2.2 Leukemia data

The goal of analyzing the leukemia data set is to classify between two types of acute leukemia, acute myeloid leukemia (AML) and acute lymphoblastic leukemia (ALL). The data set was pre-processed via RMA and contains gene expression levels produced by complementary DNA (cDNA). We first screen the genes via an independent samples t-test and select genes with p-values less than 0.05 for further analysis, resulting in 2047 genes for the final analysis.

We use a 10-fold cross-validation strategy to evaluate the performance of the different methods. The prediction error for PMMLogit, iMOMLogit, ISIS, LASSO and NP are presented in Table 4. For this analysis, PMMLogit and LASSO achieved the lowest error rate at 4.17%. The error rates of ISIS, NP, and iMOMLogit are 5.55%, 8.33%, and 19.44%, respectively. As before, when reporting the final model selected for each method, we consider analysis on the entire dataset. In the model selected by PMMLogit, there were three genes named ‘Zyxin’, ‘CD33’, and ‘ME491/CD63 antigen.’ ISIS selected two significant genes including ‘Zyxin’, ‘mRNA (clone C-2k)’, ‘CD33’, and ‘XK mRNA.’ There were two genes named ‘Zyxin’ and ‘SPTBN1’ in the model selected by iMOMLogit. LASSO found 23 significant genes in the final model while the method based on normal prior selected 943 genes in the model.

**Table 4.**
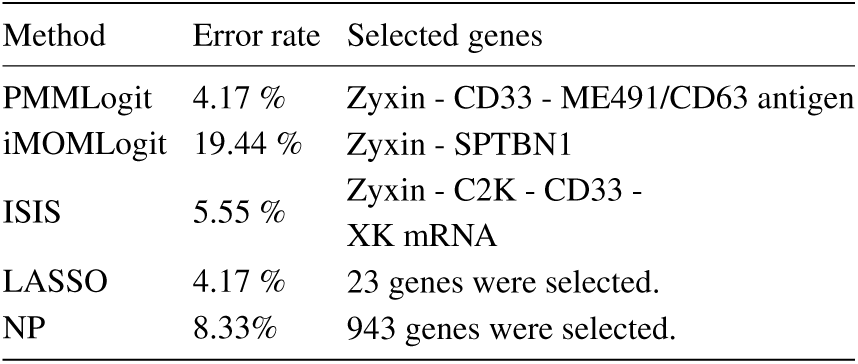
Average error rate and selected genes for the leukemia data set

*Zyxin* This gene has been reported to exert tumor suppressive functions, and mutation in acute myeloid leukemia compromises its functions. Zyxin has been listed in the most important genes reported by Golub et al. (1999) and PMMLogit selects this gene and two others in the final model. Zyxin was also included in other models, demonstrating its importance for classifying ALL and AML, which is consistent with many studies (Crone *et al*., 2011; Bernusso *et al*., 2015; Tang *et al*., 2011; Hervy *et al*., 2010; Karimi and Farrokhnia, 2014; Joshi *et al*., 2016).

*ME491/CD63 antigen* CD63 is a protein coding gene that plays a role in the regulation of cell development, activation, growth, and motility. This gene, uniquely identified by PMMLogit, was previously reported to be associated with leukemia in (Chien and Hsiao, 2013; Ben-Dor *et al*., 2000), and has been associated with early stages of melanoma tumor progression in (Hotta *et al*., 1988).

*CD33* CD33 antigen is a normal myeloid surface antigen that is expressed on the leukemic blast cells of patients with acute myeloid leukemia. This gene is selected by PMMLogit and ISIS, and has been reported in several studies (Walter *et al*., 2012; Kenderian *et al*., 2015; De Propris *et al*., 2011).

*mRNA (Clone C-2k) and XK mRNA* The gene ‘mRNA (Clone C-2k)’ is not known to be associated with leukemia and the gene ‘XK mRNA’ was only reported in one study by Au *et al*. (2005).

*SPTBN1* Spectrin is a protein that links the plasma membrane to the actin cytoskeleton, plays important role in cell structure, arrangement of transmembrane proteins, and organization of organelles. SPTBN1 (spectrin beta, non-erythrocytic 1) gene is a member of a family of beta-spectrin genes. Although SPTBN1 fusion with genes such as FLT3 are associated with atypical chronic myeloid leukemia, the SPTBN1 gene found by iMOMLogit has not previously not been shown to be directly associated with leukemia.

In summary, the genes selected by PMMLogit are supported by the literature for leukemia and colon cancer in terms of biological and statistical significance. Our proposed method also achieves the lowest prediction error rate and selects a sparse model.

## 4 Discussion

In this paper, we have constructed a Bayesian method, PMMLogit, for estimation, variable and model selection, and prediction in high-dimensional data sets with binary outcomes. We have been motivated by the challenge of selecting variables to predict binary outcomes from data sets with small sample size and a large number of predictors. We use the gold standard ‘point mass mixture’ prior, inspired by its nice theoretical properties, within a Bayesian hierarchical model, and provide its MCMC software implementation using results on mutually singular distributions provided by Gottardo and Raftery, 2008 and data augmentation for logistic regression from Polson *et al*. (2013).

Previously developed approaches in this setting have relied on the Laplace approximation or the Metropolis-Hastings algorithm for posterior computation. The first leads to inexact MCMC, and the second to potentially poor mixing. To our knowledge, the proposed method is the first implementation of a scalable and exact MCMC algorithm for logistic regression using point mass mixture priors. We perform extensive simulations to evaluate the performance of the PMMLogit approach under different sparsity and complexity situations. As expected from theory, empirical results demonstrate better performance in terms of variable selection, estimation and outcome prediction as compared to other popular models used for this task. Additionally, we have implemented the PMMLogit procedure and compared it to other methods using two real data sets, where it selects sparse models with low prediction error rates.

## Acknowledgements

The authors would like to thank James F. Dow for helpful discussions on improving the efficiency of the code for Algorithm 1 and Dr. Yelena Tarasenko for help editing the manuscript.

## Funding

The first author would like to thank the Mathematical Biosciences Institue (MBI) at Ohio State University for partially supporting this work through an Early Career Award. MBI receives its funding through the National Science Foundation grant DMS 1440386.

## Conflict of Interest

None declared.

