## Supplementary Material for "High-dimensional Bayesian phenotype classification and model selection using genomic predictors"

### Supporting Information for *'High-dimensional Bayesian phenotype classification and model selection using genomic predictors'*

Daniel F. Linder[1] and Viral Panchal[2]
[1]Medical College of Georgia, Augusta University
[2]University of North Carolina, Wilmington

#### 1 Full conditionals for Gibbs sampler

##### 1.1 Full conditonal of $\beta_k$

We derive the full conditional of $\beta_k$ componentwise. In case of the first component when $\beta_k = 0$, it follows that

$$\pi_1(\beta_k|\cdots) \propto L(y|x, \beta_0, \beta_k = 0, \beta_{-k}) \times 1$$

So, the normalizing constant is $C_1 = L(y|x, \beta_k = 0, \beta_{-k})$, where $\beta_{-k}$ is the vector of all regression coefficients except the $k^{th}$ one.

Similarly, the componentwise full conditional, $\pi_2$, is given by

$$\begin{aligned}
\pi_2(\beta_k|\cdots) &\propto L(y|x, \beta_0, \beta)\sqrt{\frac{1}{2\pi\sigma^2_{\beta_k}\tau^2_k}}\exp\Big\{-\frac{\beta_k^2}{2\sigma^2_{\beta_k}\tau^2_k}\Big\} \\
&\propto \Big[\prod_{i=1}^{n}\exp\Big(y_i - \frac{1}{2}\Big)x_i^\top\beta\int_0^\infty \exp\Big\{-\frac{(x_i^\top\beta)^2 q_i}{2}\Big\}p(q_i|1,0)\Big] \\
&\sqrt{\frac{1}{2\pi\sigma^2_{\beta_k}\tau^2_k}}\exp\Big\{-\frac{\beta_k^2}{2\sigma^2_{\beta_k}\tau^2_k}\Big\} \\
&\propto \Big[\prod_{i=1}^{n}\exp\Big\{\big(y_i - \frac{1}{2}\big)x_i^\top\beta - \frac{q_i(x_i^\top\beta)^2}{2}\Big\}\Big]\exp\Big\{-\frac{\beta_k^2}{2\sigma^2_{\beta_k}\tau^2_k}\Big\} \\
&= \Big[\prod_{i=1}^{n}\exp\Big\{k_i x_i^\top\beta - \frac{q_i(x_i^\top\beta)^2}{2}\Big\}\Big]\exp\Big\{-\frac{\beta_k^2}{2\sigma^2_{\beta_k}\tau^2_k}\Big\}, \quad \text{where } k_i = y_i - \frac{1}{2}
\end{aligned}$$

$$= \exp\Big[ -\frac{\beta_k^2}{2\sigma_{\beta_k}^2\tau_k^2} - \frac{\sum_{i=1}^n q_i {x_{ik}}^2 \beta_k^2}{2} + \sum_{i=1}^n k_i x_{ik}\beta_k - \sum_{i=1}^n q_i \sum_{\substack{j=1\\j\neq k}}^p \beta_j x_{ij} x_{ik}\beta_k \Big]$$

$$= \exp\Big[ -\frac{1}{2}\Big\{ \big((\sigma_{\beta_k}^2\tau_k^2)^{-1} + \sum_{i=1}^n q_i {x_{ik}}^2\big)\beta_k^2 - 2\big(\sum_{i=1}^n k_i x_{ik} - \sum_{i=1}^n q_i \sum_{\substack{j=1\\j\neq k}}^p \beta_j x_{ij} x_{ik}\big)\beta_k \Big\}\Big]$$

$$= \exp\Big[ -\frac{(\sigma_{\beta_k}^2\tau_k^2)^{-1} + \sum_{i=1}^n q_i {x_{ik}}^2}{2}\Big\{ \beta_k^2 - 2\frac{\sum_{i=1}^n q_i \sum_{\substack{j=1\\j\neq k}}^p \beta_j x_{ij} x_{ik}}{(\sigma_{\beta_k}^2\tau_k^2)^{-1} + \sum_{i=1}^n q_i {x_{ik}}^2}\beta_k \Big\}\Big]$$

The normalizing constant $C_2$ can be calculated as

$$\int \pi_2(\beta_k|\cdots)d\beta_k = \frac{1}{\sqrt{\sigma_{\beta_k}^2\tau_k^2\big((\sigma_{\beta_k}^2\tau_k^2)^{-1} + \sum_{i=1}^n q_i {x_{ik}}^2\big)}}\exp\Bigg[0.5\Big(\frac{\sum_{i=1}^n q_i \sum_{\substack{j=1\\j\neq k}}^p \beta_j x_{ij} x_{ik}}{(\sigma_{\beta_k}^2\tau_k^2)^{-1} + \sum_{i=1}^n q_i {x_{ik}}^2}\Big)^2\Bigg]$$

After straightforward calculation of $C_2$, the full conditional of $\beta_k$ is given by

$$(\beta_k|\cdots) \sim (1-w_k^*)\delta_0 + w_k^* N\Bigg(\frac{\sum_{i=1}^n q_i \sum_{\substack{j=1\\j\neq k}}^p \beta_j x_{ij} x_{ik}}{(\sigma_{\beta_k}^2\tau_k^2)^{-1} + \sum_{i=1}^n q_i {x_{ik}}^2}, \big((\sigma_{\beta_k}^2\tau_k^2)^{-1} + \sum_{i=1}^n q_i {x_{ik}}^2\big)^{-1}\Bigg),$$

where

$$w_k^* = 1 - \frac{1-w}{1 - w + w\sqrt{\sigma_{\beta_k}^2\tau_k^2\big((\sigma_{\beta_k}^2\tau_k^2)^{-1} + \sum_{i=1}^n q_i {x_{ik}}^2\big)}\exp\Bigg(0.5\Big(\frac{\sum_{i=1}^n q_i \sum_{\substack{j=1\\j\neq k}}^p \beta_j x_{ij} x_{ik}}{\big((\sigma_{\beta_k}^2\tau_k^2)^{-1} + \sum_{i=1}^n q_i x_{ik}^2\big)}\Big)^2\Bigg)}$$

#### 1.2 Full conditional of $\beta_0$

$$
\begin{aligned}
\pi(\beta_0|\cdots) &= L(y|x,\beta_0,\beta)\cdot\pi(\beta_0)\\
&\propto \Bigg[\prod_{i=1}^{n}\exp\Big(y_i-\frac{1}{2}\Big)x_i^\top\beta\int_0^\infty \exp\Big\{-\frac{(x_i^\top\beta)^2 q_i}{2}\Big\}p(q_i|1,0)\Bigg]\\
&\sqrt{\frac{1}{2\pi v_{b_0}}}\exp\Big\{-\frac{(\beta_0-b_0)^2}{2v_{b_0}}\Big\}\\
&\propto \Bigg[\prod_{i=1}^{n}\exp\Big\{\big(y_i-\frac{1}{2}\big)x_i^\top\beta-\frac{q_i(x_i^\top\beta)^2}{2}\Big\}\Bigg]\exp\Big\{-\frac{(\beta_0-b_0)^2}{2v_{b_0}}\Big\}\\
&= \Bigg[\prod_{i=1}^{n}\exp\Big\{k_i x_i^\top\beta-\frac{q_i(x_i^\top\beta)^2}{2}\Big\}\Bigg]\exp\Big\{-\frac{(\beta_0-b_0)^2}{2v_{b_0}}\Big\},\ \text{ where }\ k_i=y_i-\frac{1}{2}\\
&= \exp\Bigg[-\frac{\beta_0^2}{2v_{b_0}}+\frac{\beta_0 b_0}{v_{b_0}}+\sum_{i=1}^{n}k_i x_{i0}\beta_0-\frac{\sum_{i=1}^{n}q_i\big(x_{i0}\beta_0\big)^2}{2}-\sum_{i=1}^{n}q_i\sum_{\substack{k=1\\k\neq 0}}^{p}\beta_k x_{ik}^\top x_{i0}\beta_0\Bigg]\\
&= \exp\Bigg[-\frac{v_{b_0}^{-1}+\sum_{i=1}^{n}q_i{x_{i0}}^2}{2}\Big\{\beta_0^2-2\frac{\sum_{i=1}^{n}k_i x_{i0}-\sum_{i=1}^{n}q_i\sum_{\substack{k=1\\k\neq 0}}^{p}\beta_k x_{ik}x_{i0}+b_0 v_{b_0}^{-1}}{v_{b_0}^{-1}+\sum_{i=1}^{n}q_i{x_{i0}}^2}\beta_0\Big\}\Bigg]
\end{aligned}
$$

Hence, $(\beta_0|\cdots)\sim N\Bigg(\frac{\sum_{i=1}^{n}k_i x_{i0}-\sum_{i=1}^{n}q_i\sum_{\substack{k=1\\k\neq 0}}^{p}\beta_k x_{ik}x_{i0}+b_0 v_{b_0}^{-1}}{v_{b_0}^{-1}+\sum_{i=1}^{n}q_i{x_{i0}}^2},\big(v_{b_0}^{-1}+\sum_{i=1}^{n}q_i{x_{i0}}^2\big)^{-1}\Bigg)$

#### 1.3 Full conditional of $q$

$$(q_i|\cdots)\sim PG(1,|x_i^\top\beta|)$$

#### 1.4 Full conditional of $\tau_k^2$

$$
\begin{aligned}
\pi(\tau_k^2|\cdots) &\propto \pi(\beta|\sigma_{\beta_k}^2\tau_k^2)\cdot\pi(\tau_k^2)\\
&\propto \frac{1}{\sqrt{\tau_k^2}}\exp\Big\{-\frac{\beta_k^2}{2\sigma_{\beta_k}^2\tau_k^2}\Big\}\exp\Big\{-\frac{\lambda^2\tau_k^2}{2}\Big\}\\
&= \frac{1}{\sqrt{\tau_k^2}}\exp\Big\{-\frac{\beta_k^2+\lambda^2\sigma_{\beta_k}^2\tau_k^4}{2\sigma_{\beta_k^2\tau_k^2}}\Big\}\\
&\propto \frac{1}{\sqrt{\tau_k^2}}\exp\Big\{-\frac{\beta_k^2-2\beta_k\lambda\sigma_{\beta_k}\tau_k^2+\lambda^2\sigma_{\beta_k}^2\tau_k^4}{2\sigma_{\beta_k}^2\tau_k^2}\Big\}\\
&= \frac{1}{\sqrt{\tau_k^2}}\exp\Big\{-\frac{\big(\beta_k/\sigma_{\beta_k}-\lambda\tau_k^2\big)^2}{2\tau_k^2}\Big\}
\end{aligned}
$$

Let $S_k = 1/\tau_k^2 \implies 1/S_k = \tau_k^2$
So,

$$\begin{aligned}\pi(S_k) &\propto \frac{1}{S_k^2}\sqrt{S_k}\exp\Bigg\{-\frac{\Big(\frac{\beta_k}{\sigma_{\beta_k}} - \frac{\lambda}{S_k}\Big)^2}{2/S_k}\Bigg\} \\ &= S_k^{-\frac{3}{2}}\exp\Bigg\{-\frac{\lambda^2\Big(S_k - \frac{\lambda\sigma_{\beta_k}}{\beta_k}\Big)^2}{2\Big(\frac{\lambda^2\sigma_{\beta_k}^2}{\beta_k^2}\Big)S_k}\Bigg\}\end{aligned}$$

Thus, $S_k = 1/\tau_k^2 \sim Inv.Gaussian\Big(\frac{\lambda\sigma_{\beta_k}}{\beta_k}, \lambda^2\Big)$.

### 2 Additional Simulation Results

Table 1. Average FN for simulation 2.

| # of (β ≠ 0) | p | Methods | | | | |
|---|---|---|---|---|---|---|
| | | PMMLogit | iMOMLogit | ISIS | LASSO | NP |
| 5 | 301 | 1.063 | 1.060 | 4.674 | 3.444 | 92.264 |
| | 501 | 1.203 | 1.198 | 5.334 | 3.906 | 164.716 |
| | 1001 | 1.146 | 1.246 | 6.039 | 4.714 | 371.573 |
| 10 | 301 | 2.927 | 3.993 | 5.086 | 6.760 | 92.348 |
| | 501 | 3.244 | 4.228 | 5.559 | 7.667 | 168.441 |
| | 1001 | 3.414 | 4.252 | 6.096 | 8.577 | 377.105 |
| 15 | 301 | 6.319 | 8.333 | 6.302 | 10.105 | 93.464 |
| | 501 | 6.851 | 8.581 | 7.234 | 11.485 | 168.926 |
| | 1001 | 7.489 | 8.670 | 8.212 | 12.211 | 380.648 |

Table 2. Average AUC for simulation 2.

| # of ($\beta \neq 0$) | p | Methods | | | | |
|---|---|---|---|---|---|---|
| | | PMMLogit | iMOMLogit | ISIS | LASSO | NP |
| 5 | 301 | 0.820 | 0.817 | 0.751 | 0.802 | 0.675 |
| | 501 | 0.820 | 0.818 | 0.765 | 0.802 | 0.640 |
| | 1001 | 0.822 | 0.810 | 0.745 | 0.791 | 0.582 |
| 10 | 301 | 0.841 | 0.799 | 0.831 | 0.837 | 0.730 |
| | 501 | 0.827 | 0.786 | 0.823 | 0.829 | 0.681 |
| | 1001 | 0.819 | 0.784 | 0.813 | 0.810 | 0.615 |
| 15 | 301 | 0.814 | 0.750 | 0.829 | 0.851 | 0.763 |
| | 501 | 0.796 | 0.738 | 0.809 | 0.835 | 0.712 |
| | 1001 | 0.780 | 0.735 | 0.805 | 0.821 | 0.646 |

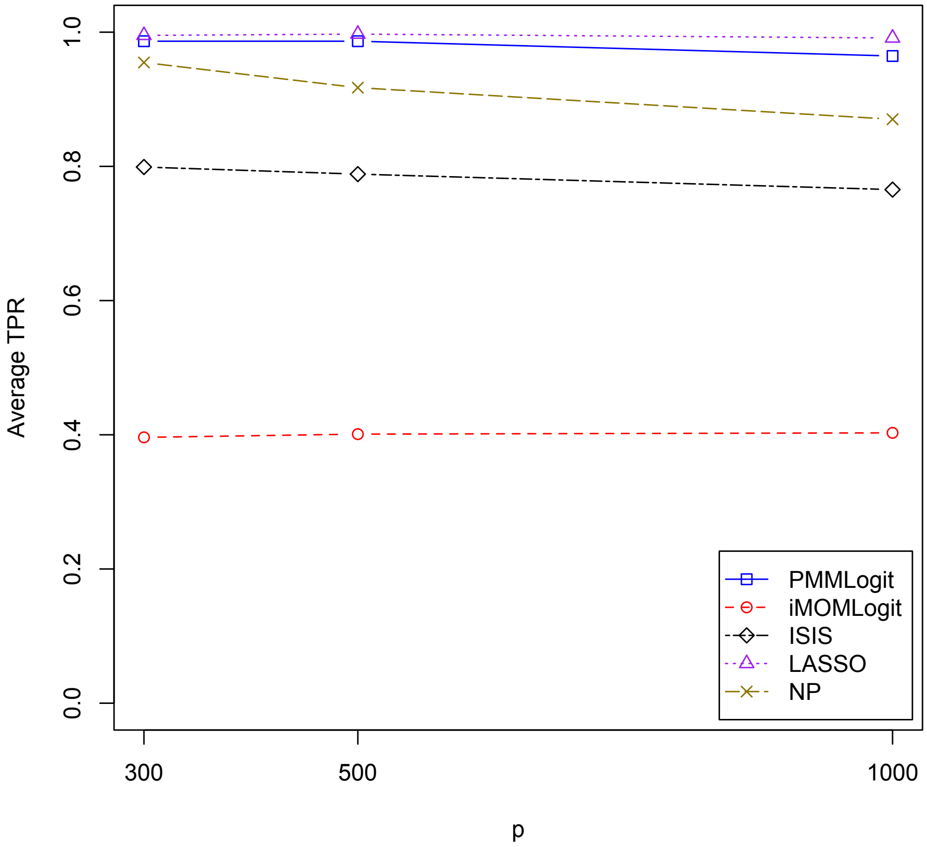


Figure 1: Average true positive rate for $\beta_5$.

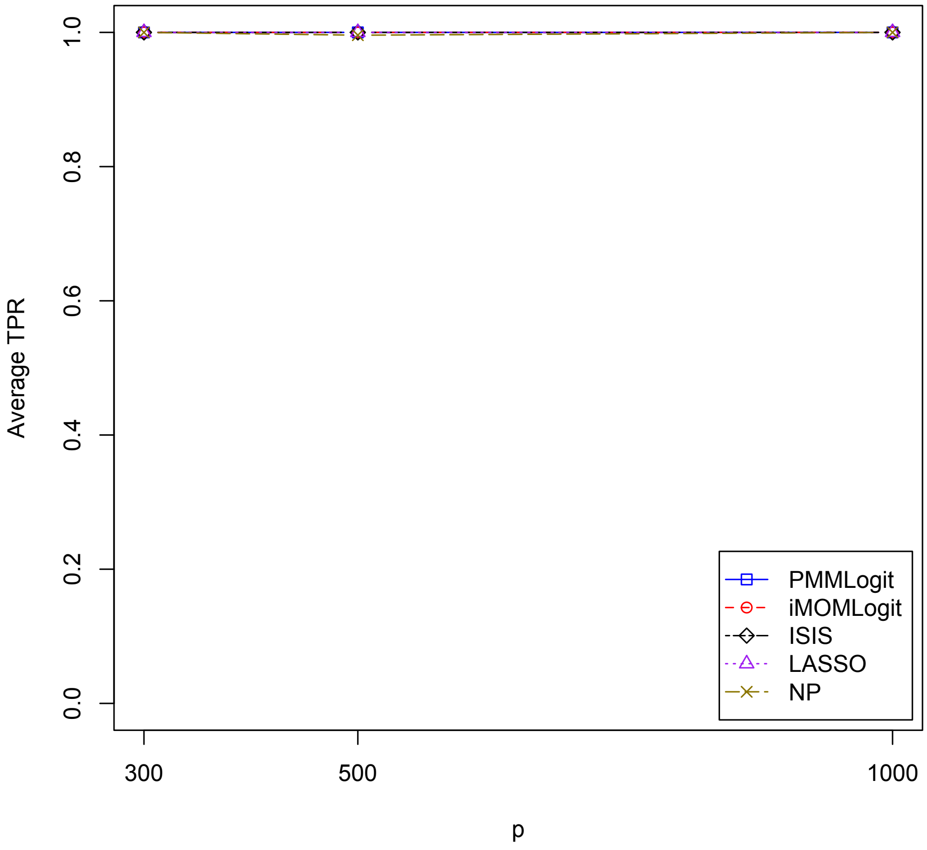


Figure 2: Average true positive rate for $\beta_1$.

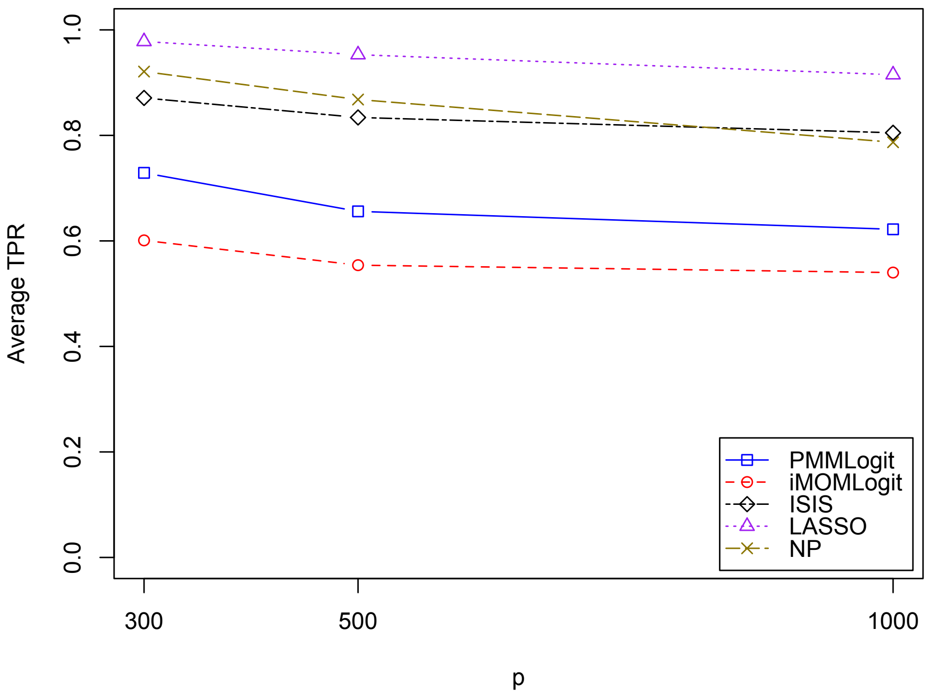


Figure 3: Average true positive rate for $\beta_4$.

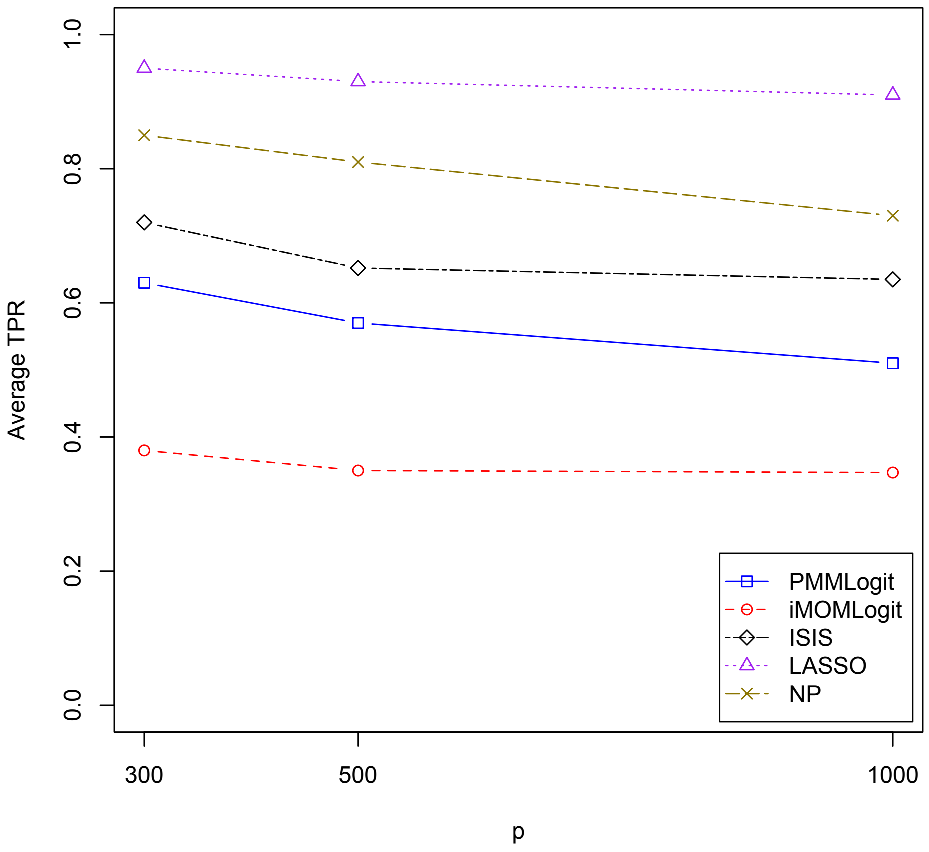


Figure 4: Average true positive rate for $\beta_6$.

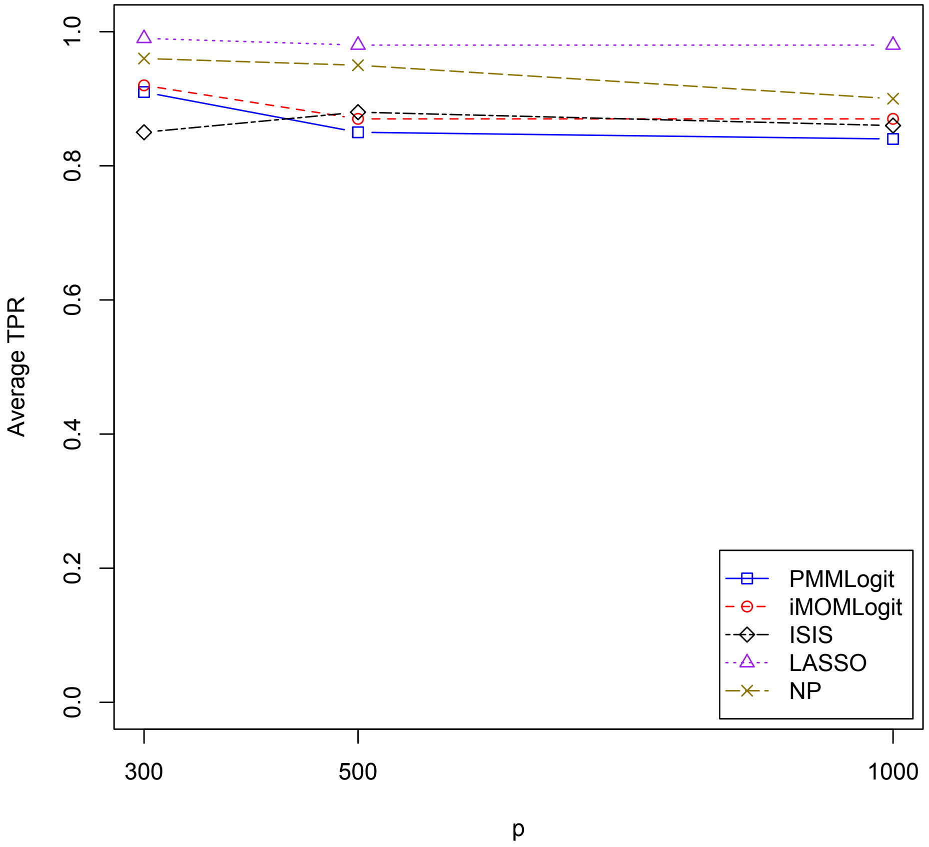


Figure 5: Average true positive rate for $\beta_2$.

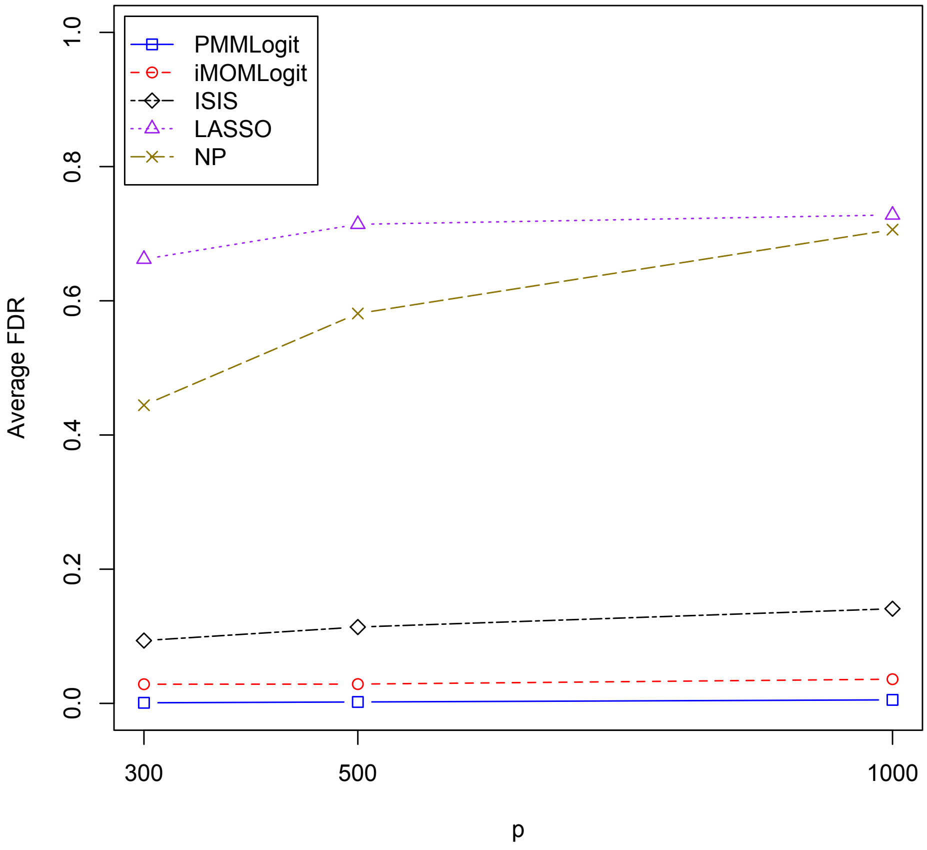


Figure 6: Average false discovery rate for $\beta_5$.

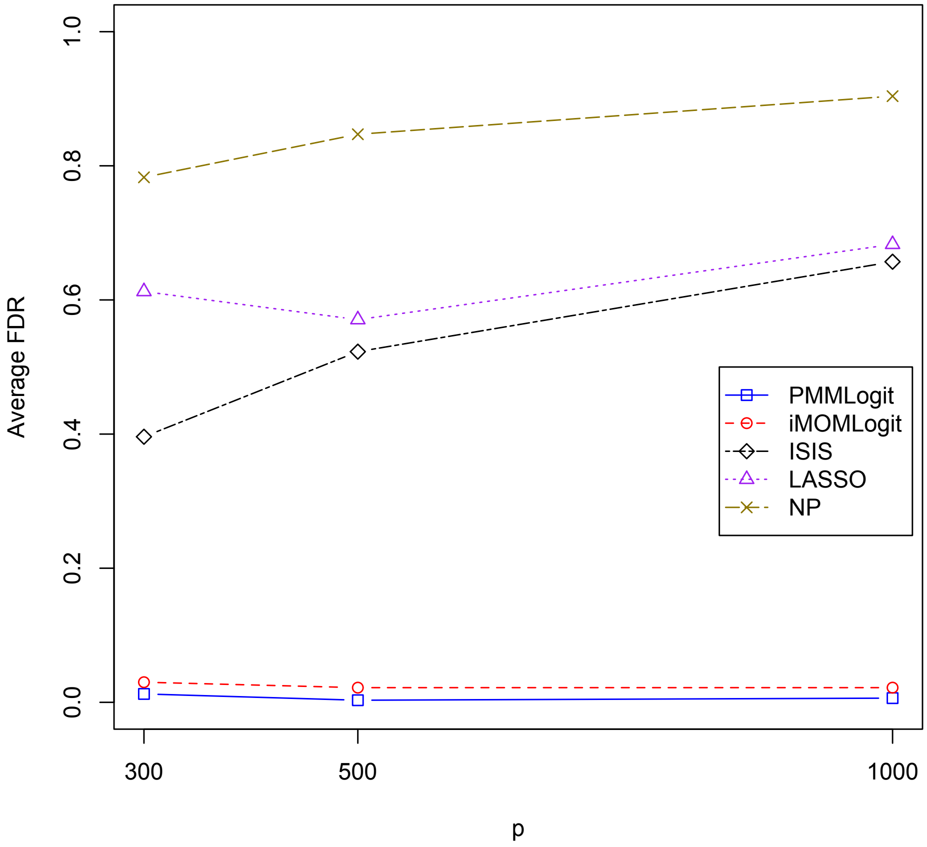


Figure 7: Average false discovery rate for $\beta_1$.

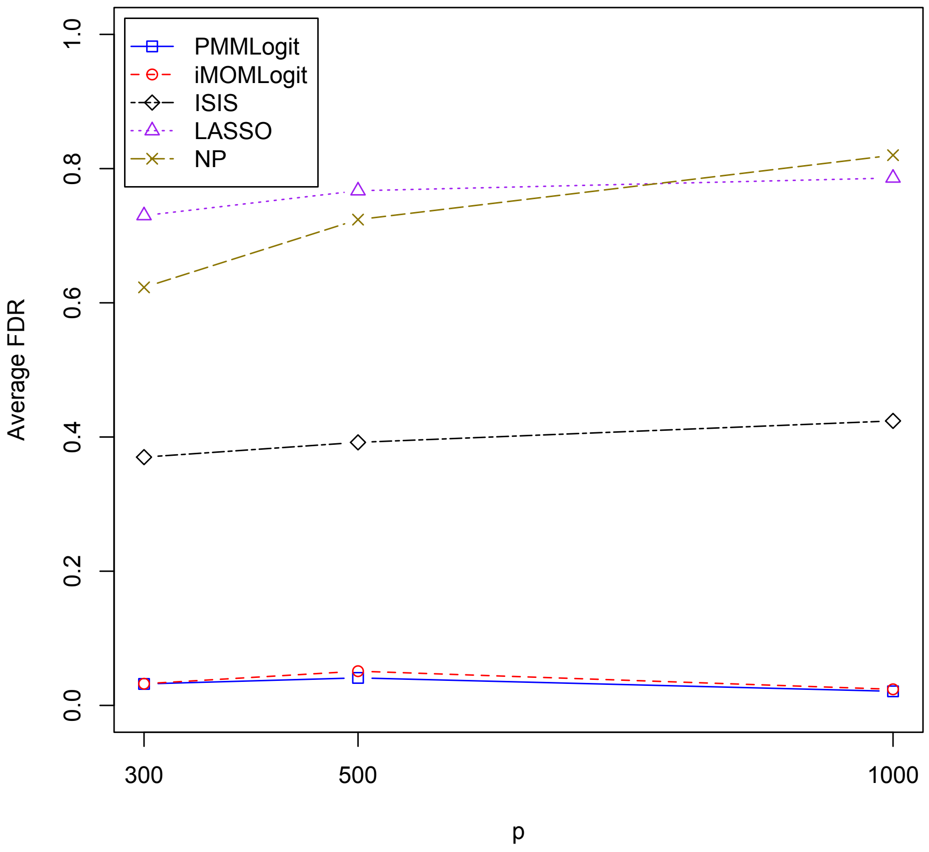


Figure 8: Average false discovery rate for $\beta_4$.

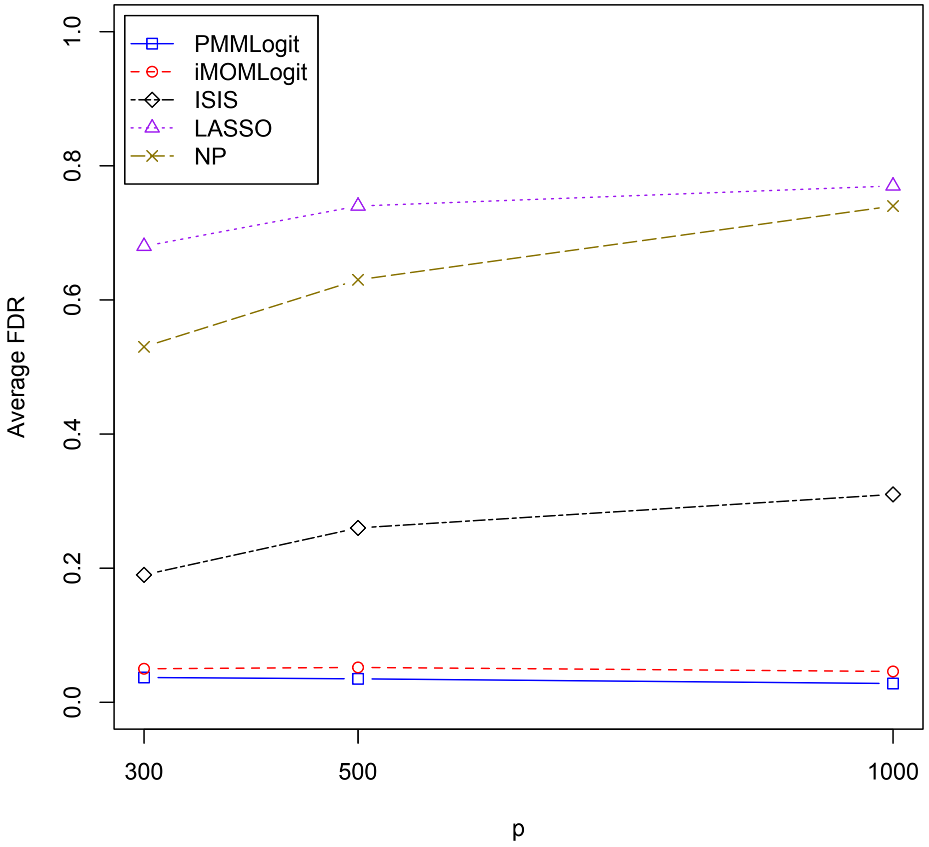


Figure 9: Average false discovery rate for$\beta_6$.

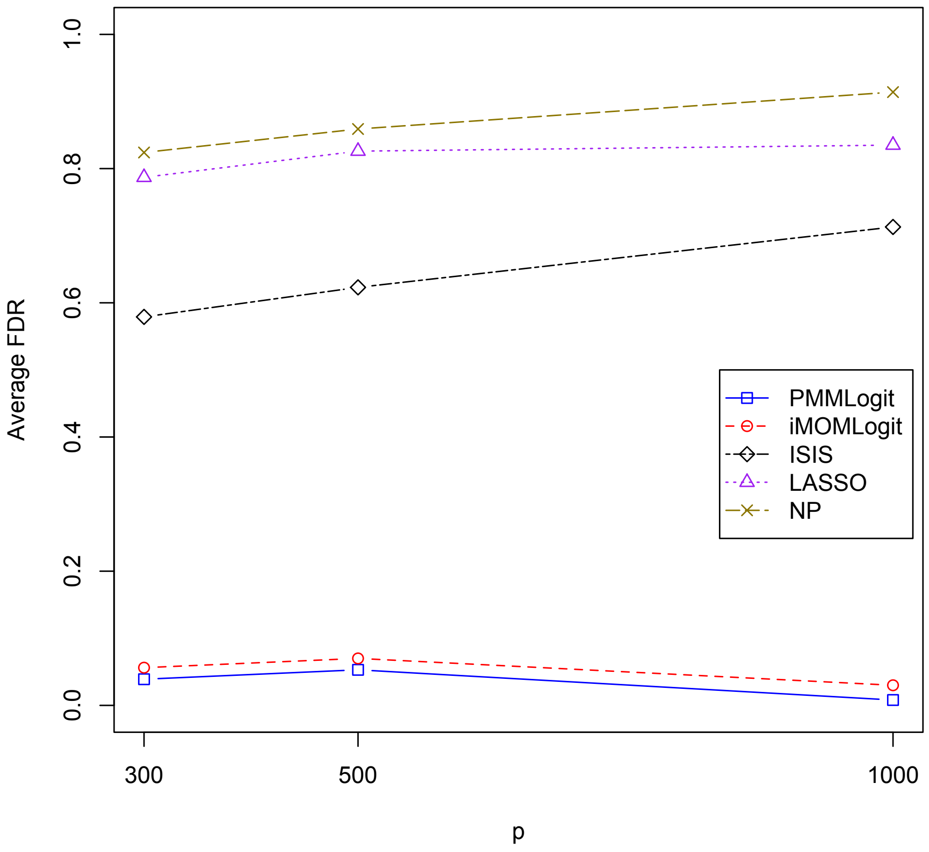


Figure 10: Average false discovery rate for $\beta_2$.
